# The Portal Project: a long-term study of a Chihuahuan desert ecosystem

**DOI:** 10.1101/332783

**Authors:** S. K. Morgan Ernest, Glenda M. Yenni, Ginger Allington, Ellen K. Bledsoe, Erica M. Christensen, Renata M. Diaz, Patricia Dumandan, Keith Geluso, Jacob R. Goheen, Qinfeng Guo, Edward Heske, Douglas Kelt, Samantha Lamb, Joan M. Meiners, Jim Munger, Raven Padgett, Carla Restrepo, Douglas A. Samson, Michele R. Schutzenhofer, Marian Skupski, Sarah R. Supp, Kate Thibault, Shawn Taylor, Ethan White, Hao Ye, Diane W. Davidson, James H. Brown, Thomas J. Valone

**Affiliations:** Department of Wildlife Ecology and Conservation, University of Florida; Biodiversity Institute, University of Florida; Department of Natural Resources and the Environment, Cornell University; School of Natural Resources and the Environment, University of Arizona; Department of Fish, Wildlife, and COnservation Ecology, New Mexico State University; Department of Biology, University of Nebraska at Kearney; Department of Natural Resource Ecology and Management, Iowa State University; Southern Research Station, U.S. Forest Service; Museum of Southwestern Biology, University of New Mexico; Department of Wildlife, Fish, and Conservation Biology, University of California Davis; Report for America; Department of Biology, Boise State University; Department of Biology, University of Puerto Rico - Rio Piedras; Sapelo Island National Estuarine Research Reserve; Division of Science and Mathematics, McKendree University; SRA International; Data Analytics Program, Denison University; National Ecological Observatory Network; Informatics Institute, University of Florida; Community for Rigor; Department of Biology, University of Utah; Department of Biology, University of New Mexico; Department of Biology, Saint Louis University; Swedish University of Agricultural Sciences; Cornell Lab of Ornithology, Cornell University; Cargill

## Abstract

This is a data paper for the Portal Project, a long-term ecological study of rodents, plants, and ants located in southeastern Arizona, U.S.A. This paper contains an overview of methods and information about the structure of the data files and the relational structure among the files. This is a living data paper and will be updated with new information as major changes or additions are made to the data. All data - along with more detailed data collection protocols and site information - is archived at: https://doi.org/10.5281/zenodo.1215988.

## Background and Summary

Long-term studies play a key role in ecology by providing unique and often foundational insights into how nature operates (Lindenmayer et al 2012; Hughes et al 2017a). Insights from long-term studies have advanced our understanding of the rapidity of species evolution (Boag and Grant 1981; Grant 1985; Arbogast et al 2006) and contributed to the development of ecological theories (Hubbell 2001) and the discovery of anthropogenic impacts on nature (Hughes et al 2017b). Despite the importance of long-term data for understanding how ecosystems and processes change over time, less than 9% of studies in ecology use data collected for more than a decade (Estes et al 2018). In ecology, data collection for a typical study spans fewer than 3 years (Tilman 1989, Estes et al 2018). Without institutional support (e.g. NSF-funded Long-Term Ecological Research sites), long-term projects can be difficult to maintain, vulnerable to both the vagaries of funding and to the longevity and interest of the scientist running it. Because long-term data is difficult to collect, there is often resistance by its collectors to making it publicly available (Mills et al 2015). Thus long-term data is highly valuable but also less available than other types of data.

This data paper describes a publicly-available, long-term study of a Chihuahuan Desert Ecosystem near Portal, Arizona in the United States (aka the Portal Project). Started in 1977, the Portal Project encompasses over 40 years of ecological research, involving both short-term and long-term experiments and monitoring of a variety of different taxa (rodents, plants, and, for many decades, ants). These data have been used in over 100 scientific publications studying competition (e.g., Munger and Brown 1981), granivory (e.g., Chen and Valone 2017), community dynamics (e.g., Ernest et al 2008), and the long-term reorganization of the ecosystem in response to habitat conversion (e.g., Brown et al 1995). Data can be downloaded from Zenodo (https://doi.org/10.5281/zenodo.1215988). The goal of this data paper is to provide an overview of the study, our available data and its structure, and the general data collection and data entry/quality assurance/quality control processes for the different data types. Detailed protocols for data collection and curation are in the metadata associated with the archived data.

## Methods

### Site Information

The Portal Project is located on a 20-hectare study site, 6 km northeast of the town of Portal, AZ. Detailed location data are provided, but site-level coordinates generally used are 31.937769, −109.08029. The project was originally established to study competition among granivores in this resource-limited system, specifically competition among rodent species, among ant species, and potentially competition between the two taxa. When the study site was established in 1977, the habitat type was best described as desert grassland habitat, though over the subsequent years it has transitioned to mixed shrubland (Brown 1998). As a desert system, it is water-limited, with the majority of precipitation occurring during two seasons. There is a short warm wet season (July - September) and another cold wet season (January - March).

Rodent and plant communities have been sampled on all plots continuously since 1977, while the ant community was sampled from 1977-2009.

Located on land managed by the Bureau of Land Management, the site is enclosed by a barbed-wire fence to exclude cattle. Inside the cattle fence are 24 experimental plots, 50 m by 50 m in size, arrayed in an approximate grid pattern (Figure 1A). Experimental plots are enclosed by a fence made of hardware cloth, about 50 cm in height and extending 20 cm underground to deter entry by burrowing rodents. Plots are designated as either controls or one of a variety of experimental manipulations, which are described below.

**Figure 1:**
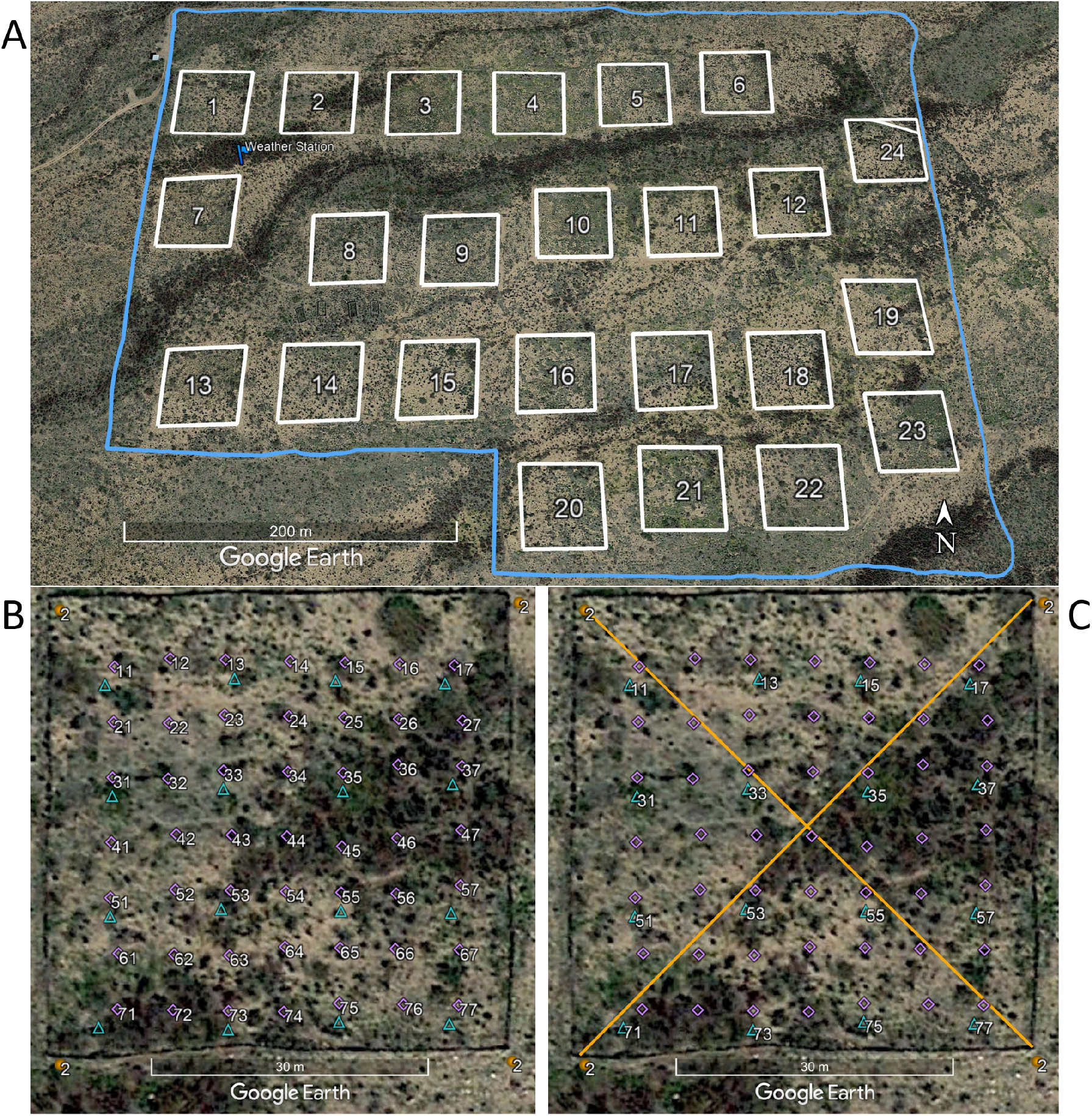
Layout of the experimental plots and sampling locations. A: Location of all 24 plots (numbered and outlined in white). Blue flag marks the weather station location. A fence to exclude cattle surrounds the entire site (blue line). The outline for plot 24 also shows how, from 1995 to 2016, the northeast corner of plot 24 was cut off, excluding 2 rodent stakes and one plant quadrat. B: Permanent rodent trapping locations (‘stakes’, purple diamonds) shown for plot 2. C: Permanent plant sampling locations. Quadrats for counting abundance (blue triangles) are interlaced with the rodent trapping locations (purple diamonds). Transects for assessing perennial cover (yellow lines) span the plot diagonally from corner to corner (yellow circles).

### Treatments

Experimental treatments applied to plots are divided into three categories: rodent access treatments, ant access treatments, and resource/seed manipulations. Each treatment category has a control state (i.e. lack of manipulation) represented in the treatment design. While half of the plots at the site have maintained relatively consistent experimental treatments focused on rodents and ants for the entire history of the study (the ‘long-term plots’), a subset of the plots have had changes to their experimental treatments during the years to focus on different scientific questions. These ‘short-term plots’ (i.e. plots with shorter length of experiments) have each been assigned to 4 different treatments over the length of the study. Details about each treatment are provided below and treatment timelines are shown in Figure 2.

**Figure 2:**
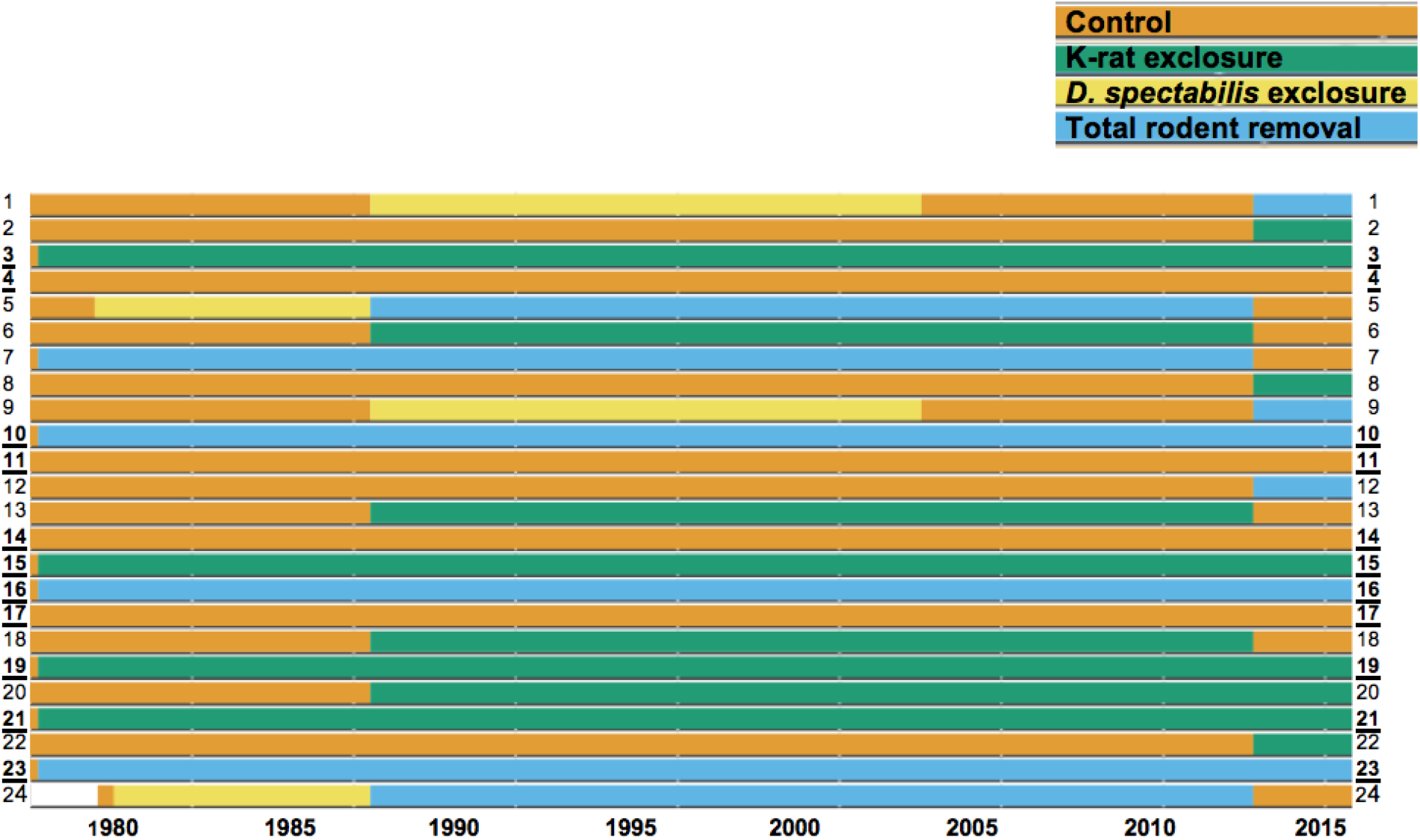
Rodent treatment assignments on plots (y-axis) over time (x-axis). All plots were controls during an initial burn-in period, before treatments were applied. Long-term plots (those which have not had their treatments altered since the original assignment) are labeled in bold and <u>underlined</u>. Plots have large gates to allow access to all rodents (“control”), medium gates to allow access to all rodents but the largest species of kangaroo rat, the Banner-tailed kangaroo rat (“*D. spectabilis* exclosure”), small gates to exclude all species of *Dipodomys* (the dominant genus) but allow access to all other rodents (“k-rat exclosure”), or no gates to exclude all species of rodent (“total rodent exclosure;” any rodents trapped on these plots are removed from the site).

#### Rodent treatments

Rodent access to plots is manipulated through 16 gates in the fence enclosing each plot. There have been 4 types of rodent treatments over the years, differentiated by the size (or absence) of the gates, determining which rodents have access to a plot. Control plots have 3.7 x 5.7 cm gates that allow access for all rodents.

Rodent exclosures have no gates in the fencing and exclude all species of rodents. Kangaroo rat exclosures have 1.9 x 1.9 cm gates that prevent species from the genus *Dipodomys* from entering. Exclosures for Banner-tailed kangaroo rats (*Dipodomys spectabilis*, the largest of the kangaroo rats), which were phased out in 2004 due to local extinction of this species, had 2.6 x 3.0 cm gates that excluded only this group. Treatments are reinforced by monthly trapping, when all rodents that should not be on that plot are removed.

Rodent treatments were changed on subsets of the short-term plots at three points in time. In January 1988, treatments were changed on 8 of the short-term plots: 2 control plots became Banner-tailed exclosures, 2 Banner-tailed exclosures became rodent exclosures, and 4 controls became kangaroo rat exclosures. After the local extinction of Banner-tailed kangaroo rats (*D. spectabilis)* from the site in the late 1990s, the plots assigned to this treatment were converted to controls in 2004. In March 2015, half of the 24 plot treatments were “flipped” to initiate a study of regime shifts: 6 controls became either rodent exclosures or kangaroo rat exclosures, 3 rodent exclosures became controls, and 3 kangaroo rat exclosures became controls. The original 11 long-term plots were kept in their long-term treatment state (see Figure 2 and *Portal_plots*.*csv* for details).

#### Ant treatments

The presence of ant species on plots was manipulated by applying a commercial poison (Mirex [Allied Chemical Corporation] until 1980, then AMDRO [American Cyanamide Company] until 2009). Poison was applied broadly across the plot (for complete ant removals) or targeted to the conspicuous mounds of *Pogonomyrmex rugosus* and *P. barbatus* (for species removal treatments). Due to a naturally decreasing abundance of *P. rugosus* and *P. barbatus* at the site over time, this species-level treatment was deemed unnecessary in 1988, and plots assigned to this treatment were converted to complete ant removals or controls. After the July 2009 sampling event, all ant treatments and ant samples were discontinued (Figure 3).

**Figure 3:**
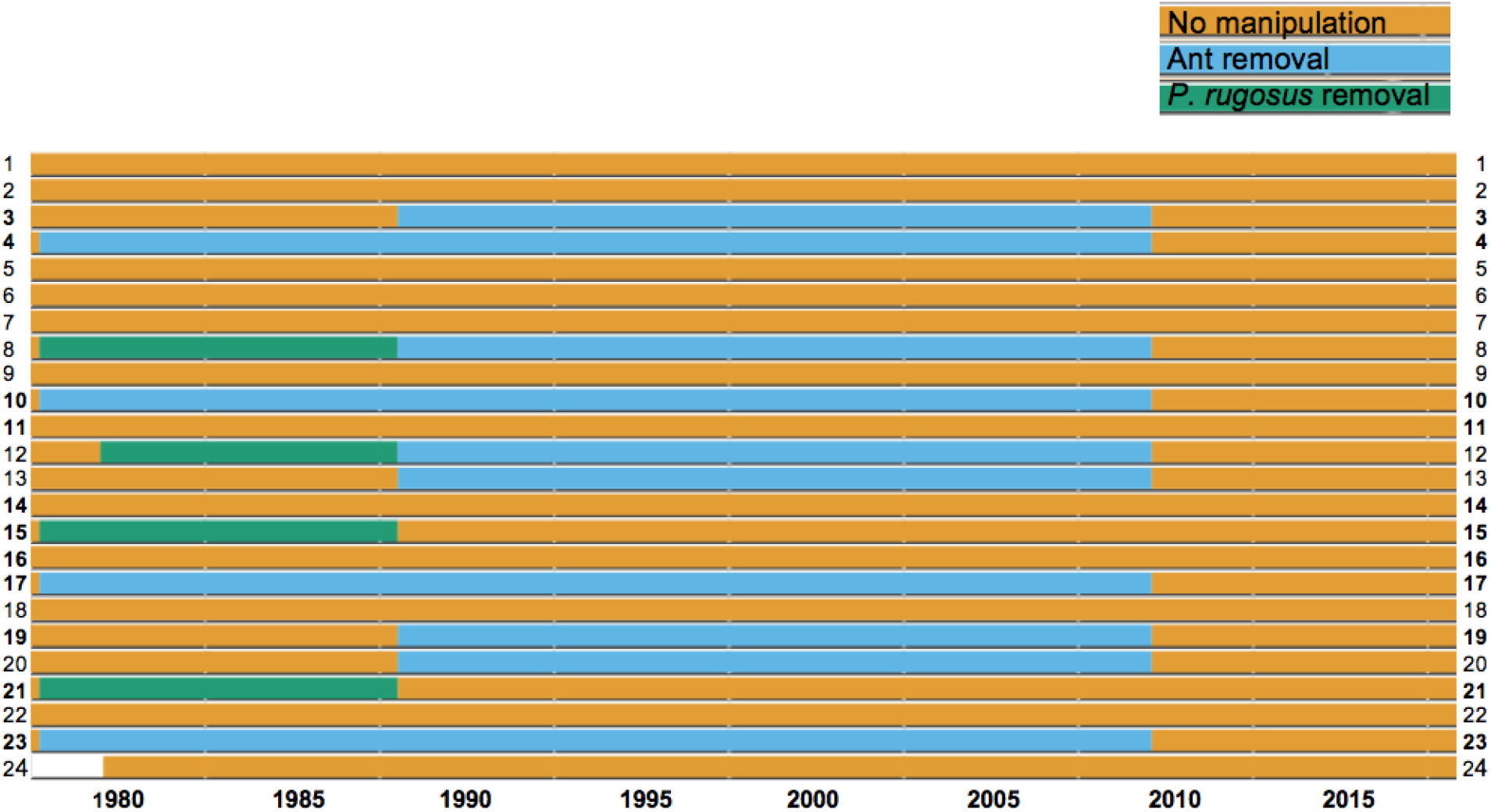
Ant treatment assignments on plots (y-axis) over time (x-axis). On non-control plots, either all ant species were removed or only the largest species of ant, *Pogonomyrmex rugosus*. Ant sampling and treatments ended in 2009.

### Resource Treatments

#### Seed addition treatments

Seed addition treatments were implemented from September 1977 to July 1985 (Brown and Munger 1985). In all treatments, 96 kg of supplemental seed (milo: *Sorghum vulgare* and/or millet: *Panicum miliaceum*) were applied to each plot per year. Treatments varied by rate of seed application: either a constant rate (equal monthly installments throughout the year) or a single yearly pulse (seed application limited to a 2-month period corresponding to peak natural seed production). Because seed size is important for how resources are used by rodent and ant consumers, this also varied by treatment: “small” seeds were created by cracking the supplemental seed to approximately ⅙ original size, and “large” seeds were the supplemental seed left whole. Eight plots were assigned to four seed addition treatments: large seeds, constant rate; small seeds, constant rate; mixed-size seeds, constant rate; and mixed-size seeds, single yearly pulse (Figure 4). In July 1985, the treatments on these plots were reassigned to plant removal, described below.

**Figure 4:**
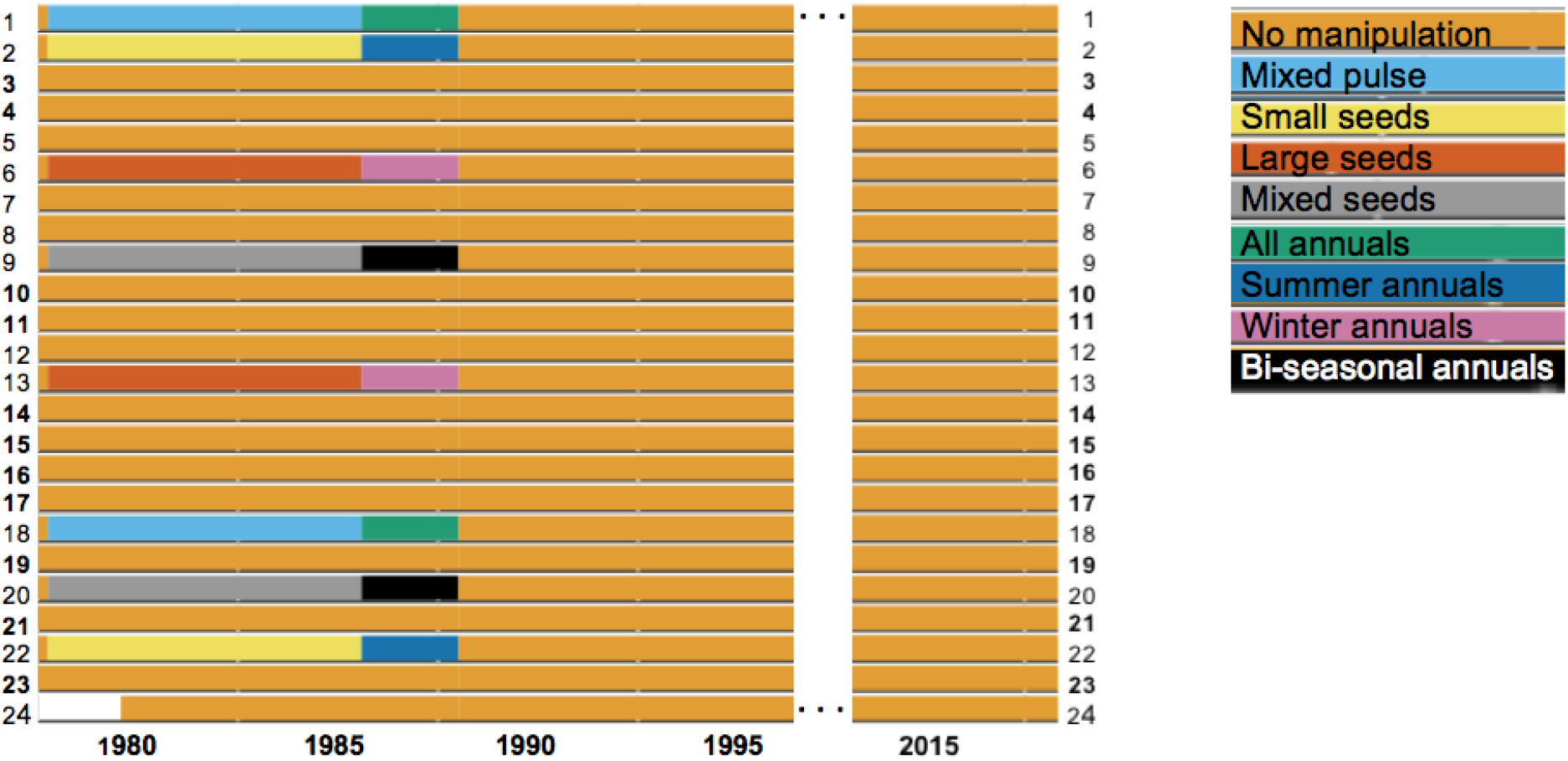
Seed additions and plant removals on plots (y-axis) over time (x-axis). All manipulations of the plant community ended in 1988. Resource treatments were generally not considered effective, or to have much impact on plot ecology.

#### Plant removal treatments

Annual plant species removal was briefly attempted by applying Roundup to assigned plots (July 1985 to December 1987). Four types of plant removal treatments, accomplished by spraying during particular seasons, targeted different groups of annual species: species which only sprout during the winter growing season (winter annuals), species which only sprout during the summer growing season (summer annuals), species which may sprout during either season (bi-seasonal annuals), or all annual species. These removals were not considered successful and were discontinued at the end of 1987. Since January 1988, there have been no direct manipulations of the plant community. Because Heske, Brown, & Guo (1993) found no significant effect of either type of manipulation to the plant community, we recommend considering these treatments no different from controls for most analyses. Nevertheless, we have included a record of their occurrence in Figure 4.

### Sampling protocols

#### Rodent sampling

We sample rodents once a month on the weekend as close as possible to the new moon. Gates related to rodent treatments are closed when traps are set in the evening and reopened the following morning. We trap half the plots (12 plots) each of two consecutive trapping nights.

Shortly before dusk, we set one Sherman trap (H. B. Sherman Traps, Inc., http://shermantraps.com), baited with millet, at each of the 49 permanent stakes in each plot (Figure 1B). We set traps within two hours of sunset and check them at sunrise the following morning. To prevent cold-induced mortality, we avoid setting traps if precipitation is expected and the temperature is below 40 F.

For captured individuals of target species, we record the plot and stake where they were caught, as well as the rodent’s species, sex, reproductive condition, weight, and hind foot length. Rodent processing is in accordance with IACUC rodent handling guidelines. Since 1991, individuals have been tagged with Passive Integrated Transponder (PIT) tags. Prior to that, individuals were identified by one or two ear tags or, occasionally, toe clipping.

Rodents caught on rodent exclosure plots and kangaroo rats caught on kangaroo rat exclosure plots are taken at least a quarter mile away and released. We record all of the routine measurements for these individuals, including any existing tags, but we do not tag untagged individuals captured on exclosure plots. Non-target species (birds, squirrels, snakes, rabbits, etc.) are identified to genus or species if possible and released immediately on site.

#### Plant sampling

Plant data has been collected once or twice annually at the site primarily in two forms: quadrats and transects. In the field, unknown species are recorded to genus if possible or with a code indicating their habit (e.g., “perennial forb”) and given a cf (“compare to”) assignment. Nearly all species observed at the site have been vouchered by the University of Arizona Herbarium. Records can be found through the Arizona-New Mexico chapter of SEINet (http://swbiodiversity.org/seinet/). We continue to voucher new species encountered at the site.

#### Quadrats

Collection of quadrat data began in 1978, though the current, higher-quality plant sampling protocol began in 1981. Sampling is conducted nearly every year, although occasional funding gaps mean samples are not fully continuous through time. Because there are two rainy seasons (January-March and July-September), there are two mostly distinct annual plant communities at the site. As such, plant sampling take place biannually. Sampling is conducted near the end of each growing season, meaning early spring for the winter plant community (typically in March or April) and early fall for the summer plant community (typically in August or September).

Each plot has 16 permanent quadrat sites (Figure 1C), marked by a rebar stake adjacent to every other rodent stage. Quadrats are placed at the 16 permanent locations by positioning the NW corner of the quadrat around the permanent rebar rebar. The northern side of the quadrat is aligned with a second permanent rebar to ensure consistent sampling. All plants rooted within the quadrats are identified and counted. Quadrat data were initially collected using 0.5 m2 quadrats but, beginning in the summer of 1981, quadrat size was reduced to 0.25 m2, 0.5 m x 0.5 m (Davidson et al 1985). In 1981-1982 only 8 of the 16 were sampled (Davidson et al 1985). As of 2015, we began visually estimating percent cover per species on each quadrat.

#### Transects

Transect data were collected every 3 years from 1989 to 2009. Each plot consisted of 4 transects (NW, NE, SW, and SE). Each transect ran from the corresponding corner of the plot (the NW transect started in the northwest corner) toward the center, each 25 m in length. Vegetation was recorded every 10 cm. This resulted in 250 sample points per transect, 1000 total points per plot. Only vegetation hits are included in the data, so there will often be fewer than 1000 records per plot. It is also possible, however, for more than one species to be recorded at the same location. There is quite a bit of variability in the recording methods used for this protocol, so these data should be used with caution. Especially if using data for all years, it is recommended to only use the transect data to calculate percent cover of shrub species. No transect data exist from 2010 through 2014.

Consistent shrub transects resumed in 2015 and are now conducted annually during each summer plant sampling event. Because the sampling protocol is slightly different, these data are housed in a separate table from the 1989 - 2009 data. Each plot has two transects: one running diagonally across the square plot from the NW corner of the plot (Figure 1C) to the SE corner and the other running from the SW corner to the NE. Only shrubs and subshrubs are recorded along the transect; annuals and herbaceous perennial species are not included. For each individual that crosses the transect, we record species identification, the beginning and end point along the transect measuring tape of where the shrub crosses the transect (anywhere in the vertical plane), and the height of the highest point of the shrub (not necessarily in the vertical plane of the transect). For 2015 only, the midpoint of where the shrub hit the vertical plane of the transect--rather than a beginning and end point--was recorded.

#### NDVI

We use a remotely sensed Normalized Difference Vegetation Index (NDVI) as a long-term measure of site-level plant productivity. Measures of NDVI are collected from 3 different types of sensors:

- Landsat (a series of satellites)
- GIMMS (an ensemble product from various AVHRR instruments on NOAA satellites)
- MODIS (one instrument aboard the Terra & Aqua satellites)

#### Phenocam

In March 2017, we installed a network camera, mounted on our weather station (location marked in Figure 1A). Approximately daily, we contribute near-surface images of our site to the PhenoCam Network (http://phenocam.sr.unh.edu/webcam/about/). The camera is north-facing and focused on Plot 2 and its surroundings.

#### Ant sampling

Between 1977 and 2009 (33 years), we monitored ants, potential competitors with rodents for seeds, on all 24 plots. This annual ant counting took place over a two-week period in July during monsoon season. Sampling recorded the species present on each plot, counted the number of colonies, estimated species abundances at crushed bait piles, or some combination of these methods depending on the year (variation detailed below).

#### Colony Data

For all diurnal ant species, we recorded the number of colonies within a 2 m radius centered around a point 2 m north of each of the 49 permanent stakes in each plot. After a colony was discovered within this search area, we identified the species present and counted the number of additional entrances within 0.5 m, which were considered to belong to the same colony. For the two *Solenopsis* species, we simply noted the presence of a colony entrance within the 2 m circle rather than count all the very numerous and tiny entrances common for this group.

For relatively rare, large-bodied ant species, colony data methodology varied slightly from the above. For *Novomessor* spp., *Pheidole desertorum, Pheidole militicida, Pogonomyrmex barbatus, Pogonomyrmex maricopa, Pogonomyrmex rugosus*, we recorded all colonies in the entire 0.25 hectare plot, noting the closest permanent stake to the colony (Davidson et al. 1985). Documentation of methods for 1984-1987 are unclear, but likely followed the above protocol.

#### Abundance Data

Ant abundance was assessed at each plot on one morning in July by counting the ants that responded to bait piles. Bait piles consisted of crushed millet placed on the ground in a 10 cm diameter circle at 25 locations in each plot. Bait piles were placed at the base of a subset of the 49 permanent rebar stakes. In rows 1, 3, 5, and 7 we placed bait at all odd column stakes (e.g. stake 11, 13, 15, 17). In rows 2, 4 and 6, we place bait piles at even numbered column stakes (e.g., stake 22, 24, 26). This created a checkerboard layout of bait piles across the entire plot. Baits were established at dawn and ants were allowed to recruit to bait piles for 1.5 hours. After 1.5 hours, all individuals of all species within a 10 cm diameter of each bait pile were counted.

### Weather monitoring

#### Site weather station

Local weather data collection has been nearly continuous at the site, beginning in 1980. The weather data spans 4 generations of weather stations, all located at approximately the same location on the site (marked in Figure 1). From 1980-1989, minimum temperature and maximum temperature were monitored with hygrothermographs, and daily values were recorded manually. Precipitation was collected with a standard rain gauge and recorded manually at irregular intervals. In 1989, an automated weather station was installed that recorded hourly temperature and precipitation. In 2002, a new station was installed that recorded hourly percent humidity, in addition to hourly temperature and precipitation. The current weather station, installed in 2016, adds several more weather measurements and calculations to the current hourly data (barometric pressure, solar radiation, evapotranspiration, wind chill, wind speed and direction).

#### External weather stations (and their integration)

Gaps in the weather data exist when a malfunction occurs in a station (e.g. the battery runs low, the station gets struck by lightning). To fill these gaps, we use data from two regional weather stations, one in the San Simon valley (US1AZCH0005), and one in the foothills near the site (USC00026716). These data are obtained from the Global Historical Climate Network Daily (http://www.ncdc.noaa.gov/ghcn-daily-description). Code for how missing precipitation values at our site are estimated from regional stations can be found in the archive.

## Data Records

In this section we describe the data files for each category of data collected at the site (i.e., plants, ants, rodents, weather) and how they relate to each other. Sub-schema for particular data types include all column names and data type (e.g., integer, character).

### Data overview

Data versions are archived on Zenodo (doi.org/10.5281/zenodo.1215988). Data are maintained live in a git repository. The repository is freely available to clone or download on Github (http://github.com/weecology/PortalData). The data contain 27 tables in 7 directories (Figure 5). Each directory contains a README file describing its tables. Additional directories contain cleaning scripts and tests used in data QA/QC (see data validation section below).

**Figure 5:**
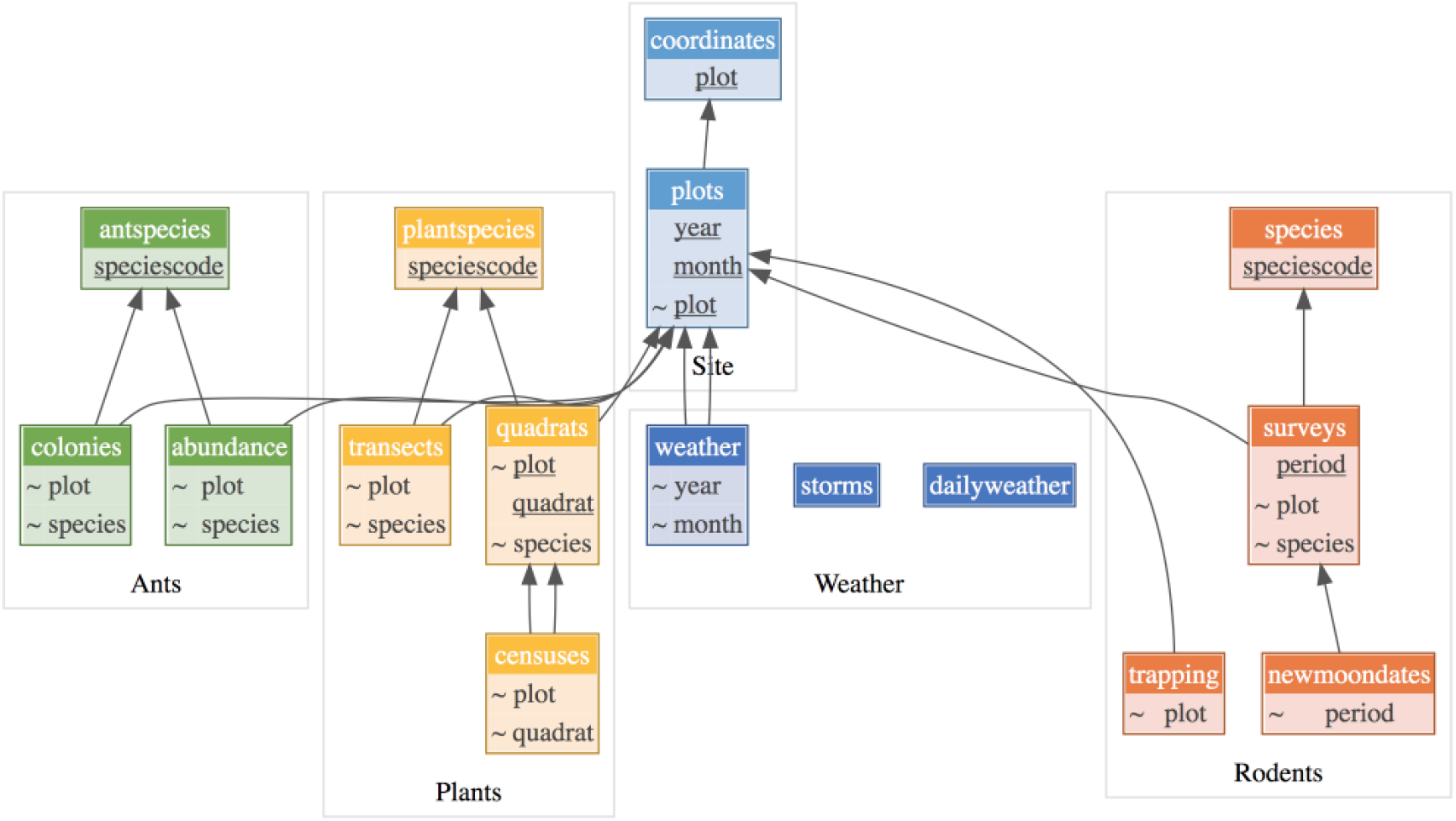
Overall database schema for entire database showing how datafiles relate to each other. <u>Underlined</u> entries denote keys or unique identifiers for a table. Entries preceded by ~ are the columns that link tables together. The name of each directory (e.g., Ants) is located at the bottom of each grey outlined box.

### Site

There are 4 site files containing plot information that is relevant to all data collected on the plots.

∘ *Methods*.*md*: A detailed description of the site and methods, including notes on data issues.
∘ *Portal_UTMCoords*.*csv*: A table of coordinates for each plot corner, rodent trapping stake, and plant quadrat, as well as the weather station location (also see Figure 1).
∘ *Portal_plot_treatments*.*csv*: Summary of the treatment history for each plot.
∘ *Portal_plots*.*csv*: A table of treatment assignment per plot over time, for the three different categories of treatment (rodent, resource, and ant). This can be used to correctly assign treatment to plot when using the raw data.

### Rodents

There are 6 rodent files: 1 data file and 5 with supporting information (Figure 6).

**Figure 6:**
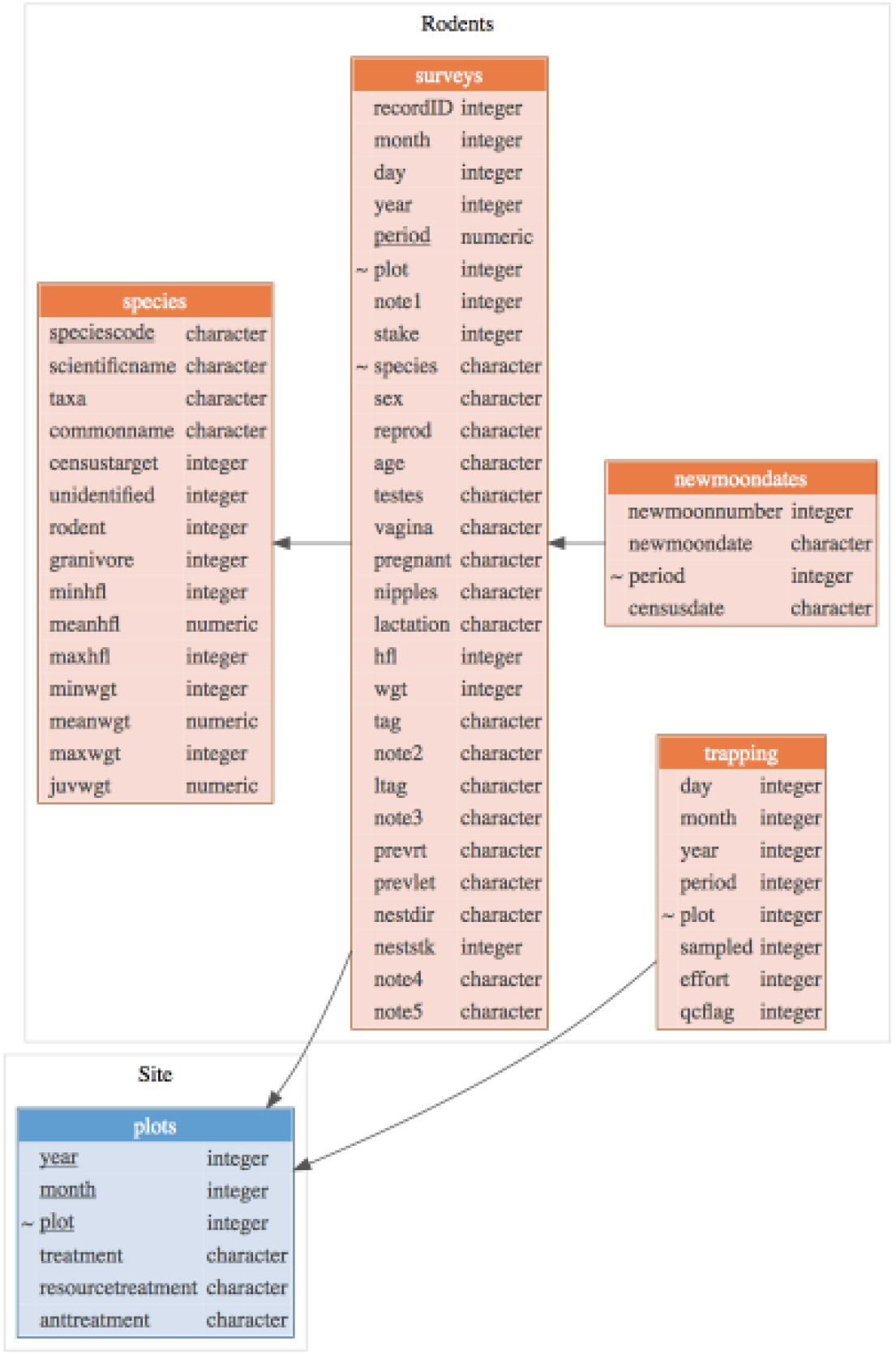
Rodent data files sub-schema. To link these data to plot treatment information, they need to be linked the relevant Site file (shown in blue). <u>Underlined</u> entries denote keys or unique identifiers for a table. Entries preceded by ~ are the columns that link tables together.

∘ *Portal_rodent*.*csv*: Contains trapping data for each individual caught, including date, period, plot, stake, tag, individual id, species code, measurements, and notes to flag common issues. Negative period codes indicate periods where there was some deviation from the usual protocols and should not be treated as regular trapping data.
∘ *Portal_rodent_datanotes*.*csv*: Key to the data flags in the “note1” column of Portal_rodent.
∘ *Portal_rodent_species*.*csv*: Key to the two-letter species codes in Portal_rodent.csv. Additional columns flag whether this species is a target species, a rodent, and/or a granivore. These columns are used to filter Portal_rodent.
∘ *Portal_rodent_trapping*.*csv*: Record of when each trapping event was done. The period codes in Portal_rodent denote the event number but do not necessarily correspond to dates. Trapping events are timed to the new moon as much as possible. Occasionally, months are skipped due to weather or other extenuating circumstances.
∘ *Portal_suspect_stakes*.*csv*: A record of non-standard stake values. For normal trapping events, these values may be misrecorded or incomplete (i.e. only the row or column is known, or there were multiple records for a single stake). Off-plot trapping data will also have non-standard stake values, but these data should not be combined with regular trapping data. *moon_dates*.*csv*: A time series of new moons, used to translate between period numbers and new moon dates

### Plants

The plant data are contained in 6 files: 2 data files and 4 files with supporting information that help with using the data files effectively (Figure 7).

**Figure 7:**
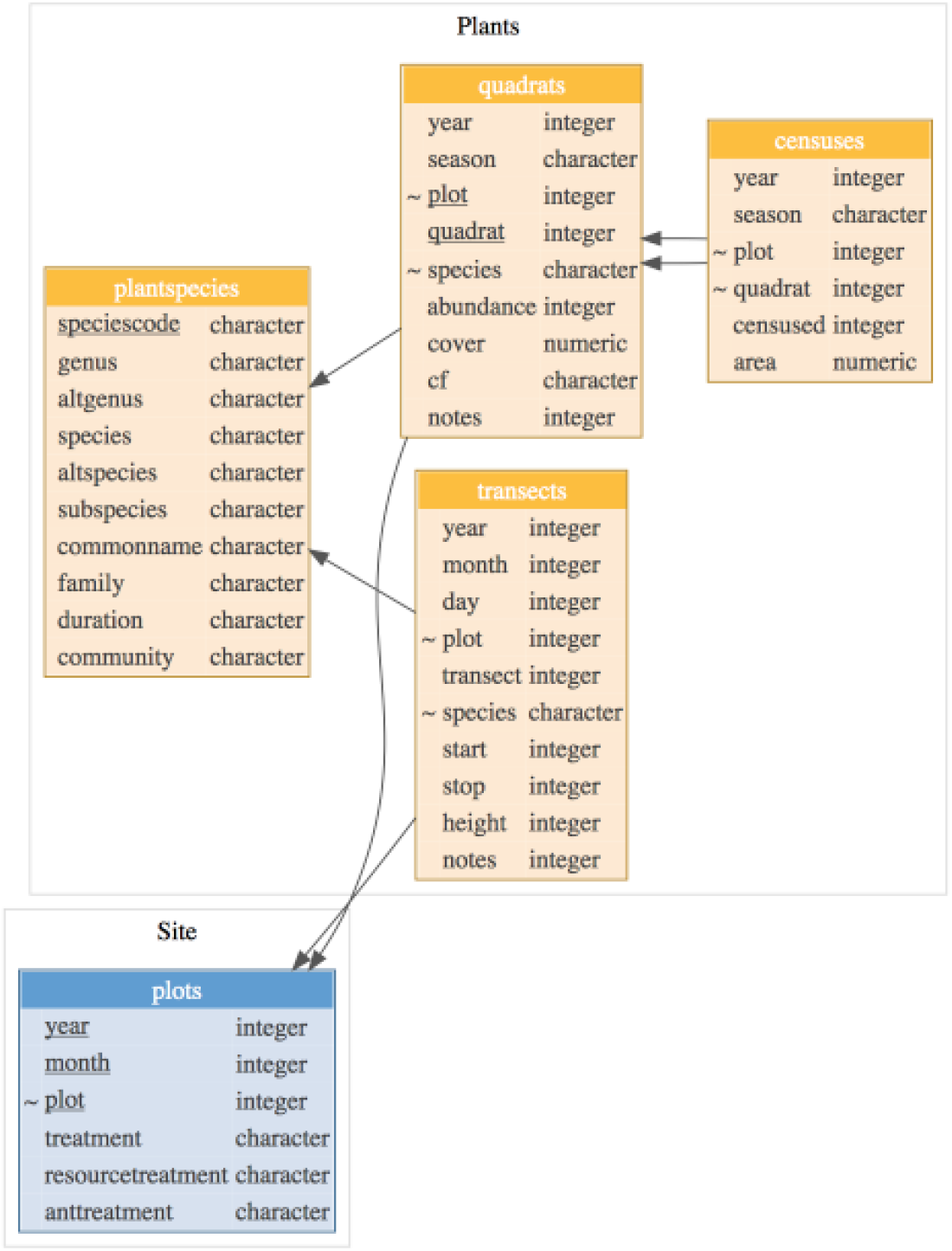
Plant sub-schema.To link these data to plot treatment information, they need to be linked with the relevant Site file (shown in blue). <u>Underlined</u> entries denote keys or unique identifiers for a table. Entries preceded by ~ are the columns that link tables together.

- *Portal_plant_census_dates*.*csv*: table including data on whether sample events occurred during summer and/or winter for each year. It also includes start and end dates if known.
- *Portal_plant_censuses*.*csv*: table including data for each conducted sampling event on whether or not each quadrat was surveyed and, if so, the quadrat area used for that sample. This can be used to differentiate between real zeros and missing data.
- *Portal_plant_datanotes*.*txt*: text file with descriptions of each data note included in all plant data files
- *Portal_plant_quadrats*.*csv*: table including species identifications and abundance for each species found in each quadrat. Starting in 2015, it also includes an estimate of percent cover.
- *Portal_plant_species*.*csv*: table with the plant species codes used in both the quadrat and shrub transect data tables, as well as information about each plant, including: family, genus, species, previous Latin binomial information, common name, community (e.g. shrub, summer annual, winter annual) and duration (annual/perennial).
- *Portal_plant_transects_1989_2009*.*csv*: table of shrub transect data, including species identification, transect location, and point on transect.
- *Portal_plant_transects_2015_present*.*csv*: table of shrub transect data, including species identification, start and stop points, and maximum height.

### NDVI

NDVI consists of a single raw data table. LANDSAT provides the longest most consistent time series. However, LANDSAT sensors have changed over time (the data start with using Landsat 5 and currently data are obtained from LANDSAT 9). The README contains instructions for how to make adjustments between sensors to get one consistent time series but these adjustments are made automatically in the portalr package.

- *ndvi*.*csv:* contains raw NDVI values for all sensors at all available dates
- *monthly_NDVI*.*csv*: is retained for backwards compatibility but no longer used. It contains monthly NDVI values for July, 1981 to December, 2013.

### Phenocam

Images are publically available via the PhenoCam Network (https://phenocam.sr.unh.edu/webcam/sites/portal/).

### Ants

There are 4 ant files: 2 data files and 2 files with supporting information (Figure 8).

**Figure 8:**
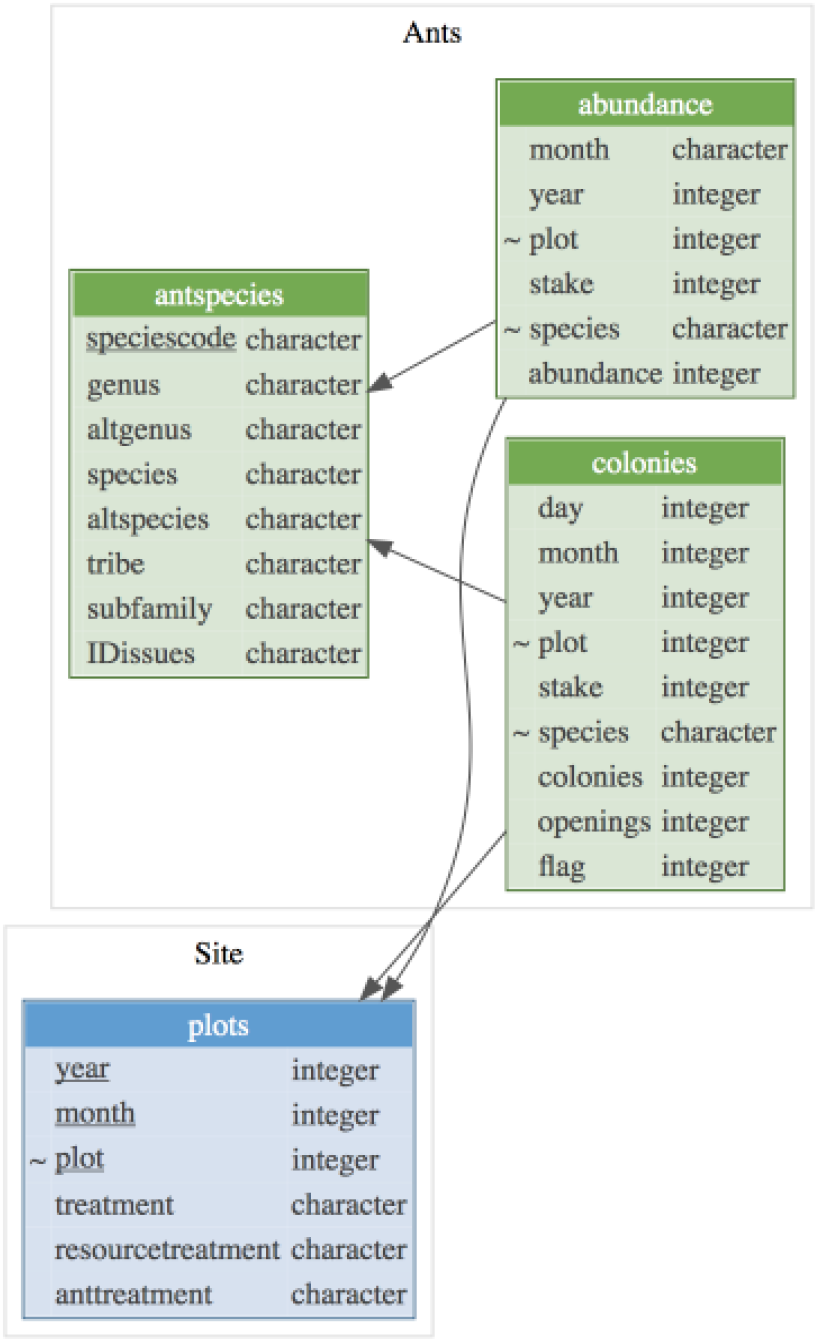
Ant sub-schema. To link these data to plot treatment information, they need to be linked with the relevant Site file (shown in blue). <u>Underlined</u> entries denote keys or unique identifiers for a table. Entries preceded by ~ are the columns that link tables together.

**Figure 9:**
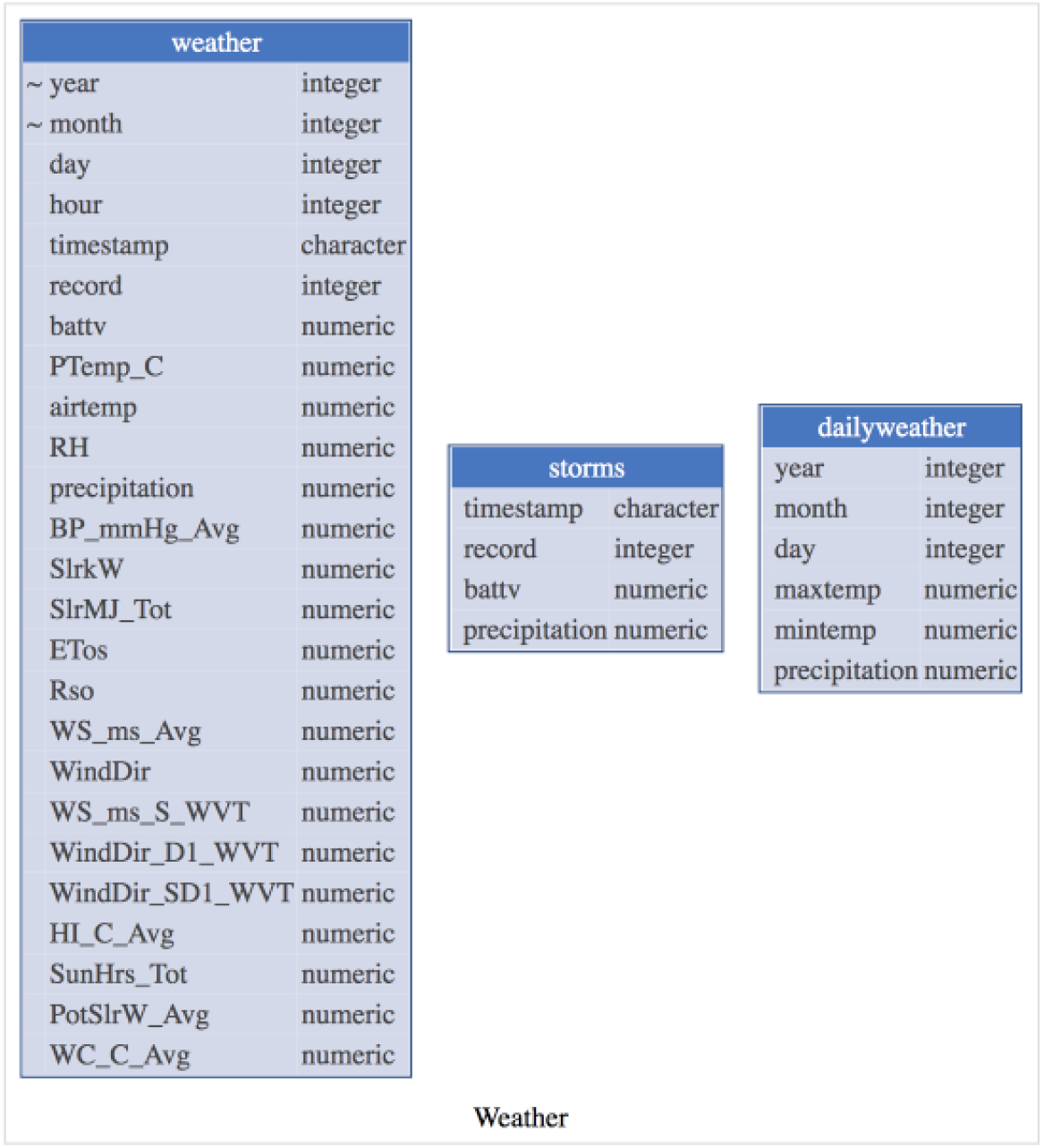
Weather sub-schema. Entries preceded by ~ are the columns that link tables together.

∘ *Portal_ant_bait*.*csv*: Raw validated data from the ant bait sampling protocol, a time series of abundances.
∘ *Portal_ant_colony*.*csv*: Raw validated data from the ant colony sampling protocol, a time series of colonies and colony openings.
∘ *Portal_ant_dataflags*.*csv*: Guide to the ant dataflags used in the flag column of Portal_ant_colony.csv.
∘ *Portal_ant_species*.*csv*: Species codes used in Portal_ant_bait.csv and Portal_ant_colony.csv.

### Weather

There are 6 weather files, all containing data.

∘ *Portal_weather*.*csv*: Hourly weather recorded from the site weather station 1989-present.
∘ *Portal_weather_19801989*.*csv*: Approximately daily weather manually recorded from 1980 to 1989.
∘ *Portal_storms*.*csv*: Precipitation (mm) data is recorded into this table every 5 minutes during a precipitation event. Provides finer-scale storm data than what is available in Portal_weather.csv.
∘ *Portal_weather_overlap*.*csv*: Two stations were in operation from 2017 - 2018. This table includes data from both stations, to use for monitoring the calibration of the sensors.
∘ *Sansimon_regional_weather*.*csv*: Daily weather from a station in the San Simon valley, used to fill gaps when the Portal site station is down.
∘ *Portal4sw_regional_weather*.*csv:* Daily weather from a station in the foothills near the town of Portal, AZ, used to fill gaps when the Portal site station is down.

## Data Validation

### Overview

Core data tables (raw data collected in the field) are validated before appending to the dataset. After QA/QC (quality assurance/quality control) of the core data tables, supplementary tables are automatically generated from the core data tables. In general, data QA/QC follows three steps: manual data entry and cleaning, automated data testing, and automated updating of supplementary tables. See Yenni et al. (2018) for an in-depth explanation of our data management procedures.

Data are first double-entered in an Excel spreadsheet with restricted fields to prevent typos. An R script is used to check for discrepancies between the double-entered versions, to further identify and remove typos, and to run a series of data validation checks. For all data types, the data validation checks ensure that all records are consistent with allowable values (i.e., realistic values given how the data are collected) and previous records (i.e., repeat observations of a single individual should have the same species). Further specifics of data validation for each data type is described below.

Following data validation, the data are appended to the database. A series of additional checks runs automatically to ensure that the new version of the data meets expected quality standards. This also triggers additional automated scripts to update the supplementary data tables associated with that data type. Below, we explain the specifics of these tables for each data type.

### Rodents

New rodent data must be consistent with realistic values for each rodent species, the possible stake values, and any previous records of specific individuals. For example, species-specific tests flag any records with unusually high or low hind foot or weight measurements, and datasheet values for these measurements must be manually confirmed. If a stake value occurs for two records in a given plot, those records are flagged and must be manually investigated. If a new record for an individual conflicts with a previous record regarding the individual’s identification, the records must be manually investigated and the user must decide whether to correct one of the records or flag the individual for closer examination in future trapping events.

After new records have been error checked, all rodents are assigned a unique ID that can be used to track individuals over time. Due to the length of the dataset, ear tag numbers and occasionally even PIT tag sequences get repeated, and therefore cannot be used directly as an individual ID. The unique ID controls for repeated tags in different individuals and individual recapture in the same trapping session We assume that a repeated tag from a new individual if > 3 years occurs since the previous record of the tag number in our database. There are also many individuals that do not get tagged at all (e.g. removals). Though these likely include some recaptures, the unique ID makes the simplest assumption that each untagged animal captured on a removal plot is a new individual. Once the new rodent records are cleaned and appended to *Portal_rodent*.*csv, Portal_rodent_trapping*.*csv* and *Portal_plots*.*csv* automatically update to include the dates and treatments for the new trapping event. Finally, these updated tables also undergo automated quality checks.

### Plants

#### Plant Quadrats

Plant quadrat data are checked to ensure that no quadrats are missing or counted twice and for valid species, abundance, cover, and cf values. If necessary, new species are manually added to the *Portal_plant_species*.*csv* table. Following the addition of new data, automated data quality tests are run, the *Portal_plant_census_dates*.*csv* and *Portal_plant_censuses*.*csv* tables are updated to include the new sample event, and they are re-tested, as well.

#### Shrub Transects

Shrub transects are checked to ensure that no transects are missing or counted twice and for valid plot, transect, species, start, stop, height, and cf values. If necessary, new species are added to the *Portal_plant_species*.*csv* table. Following the addition of these data, the automated data quality tests are re-run.

#### NDVI

Pinzon & Tucker (2014) performed a comprehensive calibration to correct for anomalies arising from different sensors, orbital drift, volcanic eruptions etc. When a pixel did not meet quality standards, it was interpolated with splines from nearby values in time. If nearby values were not available, it was set to the seasonal average for that pixel. We kept only NDVI values which had a QA value of 1, 2, or 3. These indicate “good” values (1 and 2) or values interpolated from splines (3).

#### Ants

Because ant data collection is no longer active, the data validation process happened once, for the original digital ant data entry, but does not continue on a regular basis. Data were tested for valid species, valid colonies and openings values (colony data), and valid abundance values (abundance data).

#### Weather

Unlike the above data, weather data collection is entirely automated, as is its testing. Hourly values are remotely collected from the weather station web address, automatically tested for validity, and appended to the weather table.

## Usage Notes

### Data usage recommendations

When using the data to conduct analyses that rely on or assume that sampling is consistent over some period of time, there are important issues with the data to consider.

1. Many plots shift their treatments over the course of the study. If the user is interested in analysing specific treatments, they should check which years those treatments were in effect (Figure 2; Table: *Portal_plots*.*csv*) and constrain their analyses accordingly.
2. While plant and ant sampling events were either completed or not completed, rodent trapping is sometimes only partially completed. Partial rodent trapping occurs when weather or travel complications result in only one night of trapping being conducted. Only half of the plots are trapped during these partial events. Information on which rodent samples were complete or partial events can be found in Table: *Portal_rodent_trapping*.*csv*.
3. Rodent sampling events that do not conform to protocol are indicated with a negative period code. It is recommended that these periods be excluded before analysis.
4. In the early years of the study, there were changes in the number of quadrats counted per plot for each season of plant data. In 1981 and 1982, only 8 quadrats were surveyed on each plot in each season (8 in winter and the other 8 in summer). Sampling protocols are consistent starting in 1983 (all 16 quadrats counted in every season). If using data before 1983, users should correct for these shifts in sampling effort.
5. Plant quadrats were originally established to avoid the inclusion of shrubs and perennial species. Additionally because of this focus on annuals, prior to 1989, perennial species were not counted on plant quadrats. Since 1989, all species occurring in a quadrat were recorded, and perennial species have of course begun to encroach upon the permanent quadrats. If using the data before 1989, care should be taken to account for the bias against perennial species.
6. Recording methods for the 1989 - 2009 transect data are highly variable. Analyses of these data should only be done after thoroughly reading the methods and examining the data. One should not assume these data include all species seen on a transect. Years vary in what species were counted during transect sampling. Some amount of species filtering should be done before analyzing the data.In most cases, you will only want to use the shrub species. Most years have a transect and point value, recording the location on the plot for the individual plant. 2004 and 2009 have no record of these locations. The data were collected under the same protocol, however, so they can still be used to calculate percent cover in the same way.

### R Package

An R package is being developed by the research group to help simplify access and processing of data for analysis. The portalr package is not formally released yet, but can be installed directly from GitHub through devtools (devtools::install_github(“weecology/portalr”)) or cloned from the GitHub repository (http://github.com/weecology/portalr). Documentation on the package is also available in the portalr repository.

### Data versions

Every time new data is added or changed in the PortalData repository on GitHub, it triggers a new archived release of the data on Zenodo. Each archived version has a unique data version number to allow easy reference to a particular state of the data (e.g., 1.17.0) and follows general semantic versioning protocols. A change in the patch version (i.e., 1.17.0 to 1.17.1) denotes a fix to data already in the database. A change in the minor version (i.e., the 1.17.0 to 1.18.0) denotes an addition of new data. Finally, a change in the major version (i.e., the 1.17.0 to 2.0) denotes a major structural change in the database that we anticipate will break back-compatibility of the data with existing analysis code (i.e. if you change to that version of the data, any code you already made for working with the data may stop working). All previous versions of the data are available through the DOI for the data archive: 10.5281/zenodo.1215988.

## Acknowledgements

A long-term study is not possible without support from funding agencies, the scientific community, and the friends and families of the scientists involved with the project. The Portal Project has been lucky enough to be nearly continuously funded by the National Science Foundation, most recently by grant #2430620 (https://www.nsf.gov/awardsearch/show-award?AWD_ID=2430620). The Department of Energy also provided funding for the site through a grant early in the study. R. Diaz was supported by an NSF Graduate Research Fellowship (DGE-1315138). Many people have helped with data collection over the years. Friends, family, and graduate students helping each other with their research projects have all helped out at the site. For over 20 years, Alan Ernest helped with the monthly rodent trapping, providing incalculable assistance.

## Notes

https://github.com/weecology/PortalData

### Competing Interest Statement

The authors have declared no competing interest.

### Summary of Updates

New version adds info on unique rodent IDs. Adjusts NDVI values for LANDSAT satellite changes. Updates author list, adding new authors responsible for data collection and updates institutions for other authors. Updates NSF grant number.

